# Multimodal Profiling of Repair-Associated Immune Dynamics in a Mouse Model of Menstruation

**DOI:** 10.64898/2026.02.06.704418

**Authors:** Rebecca J. Ainslie, Phoebe M. Kirkwood, Hanning Li, Symeon Gerasimou, Philippa T.K. Saunders, Ioannis Simitsidellis, Douglas A. Gibson

**Affiliations:** Centre for Reproductive Health, Institute for Regeneration and Repair, University of Edinburgh, Edinburgh BioQuarter, 4-5 Little France Crescent, Edinburgh EH16 4UU; Centre for Inflammation Research, Institute for Regeneration and Repair, University of Edinburgh, Edinburgh BioQuarter, 4-5 Little France Crescent, Edinburgh EH16 4UU

## Abstract

Despite the importance of inflammation to menstruation, we lack detailed understanding of how immune dynamics contribute to endometrial repair. We performed detailed phenotypic characterisation of uterine immune cells using single cell RNA sequencing, flow cytometry and multiplex immunohistochemistry in a mouse model of simulated menstruation. Uterine tissues were collected from age-matched controls or from key phases of menstruation, including tissue breakdown, repair and remodelling. Our findings reveal distinct compositional changes across different phases of menstruation and highlight predominant roles for monocytes, macrophages, and neutrophils in endometrial repair. Immunohistochemistry revealed the spatial association of these myeloid cell subsets with areas of tissue repair and remodelling. Bioinformatic analysis highlighted key roles for monocyte, macrophage and neutrophil signalling during endometrial repair with thrombospondin 1 and secreted phosphoprotein 1 emerging as key signalling pathways. These data significantly advance our understanding of menstrual physiology and identify potential therapeutic targets for menstrual disorders.

**summary:** This study investigates immune cell dynamics in a mouse model of menstruation, highlighting roles for monocytes, macrophages, and neutrophils in non-fibrotic endometrial repair, and identifies key signalling pathways as potential therapeutic targets for menstrual disorders.

## Introduction

Menstruation is characterised by an inflammatory cascade resulting in the breakdown and shedding of the functional layer of the endometrium followed by phases of endometrial repair and remodelling [1–4].This extensive inflammatory process must be tightly regulated to maintain reproductive function, with deficits in endometrial repair associated with reproductive health disorders [5]. Despite the importance of inflammation to menstruation and reproductive health, our current understanding of how inflammatory cells contribute to endometrial repair is limited.

Studies investigating endometrial immune cell composition during human menstruation have relied on immunohistochemistry analysis of fixed tissues with limited direct profiling of immune cell subsets. These studies have reported that the abundance of monocytes and macrophages dramatically increase prior to menstruation [6] whilst neutrophils are increased during endometrial breakdown and repair [7, 8]. Profiling of lymphoid cells is more limited. A study in the 1990’s suggested CD8+ T cells are increased post-menstruation during the early-mid proliferative phase of the cycle [9]. B cells are sparse in the endometrium throughout the cycle and therefore their role during menstruation has not been investigated [10]. In contrast, while uterine natural killer cells (uNKs) are abundant in the late secretory phase and in association with fertility regulation, their numbers are reportedly low during menstruation [11, 12] suggesting they may not play a significant role in endometrial repair.

The generation of human scRNAseq datasets have increased our understanding of the composition of endometrial immune cells in the proliferative and secretory phases of the menstrual cycle [13, 14]. However, studies profiling immune cells in tissue samples from the menstrual phase are lacking. To overcome these limitations we, and others, have developed and validated a refined mouse model of simulated menstruation [15] to gain insights into tissue dynamics, immune cell composition and mechanisms of scarless repair. Although mice don’t naturally menstruate, the physiological changes that occur in the human endometrium can be modelled using hormonal priming and artificial stimulation of implantation. Following hormone withdrawal, uterine tissues undergo reproducible breakdown and repair, recapitulating key features of human menstruation including rapid re-epithelialisation, stromal cell remodelling and endothelial cell proliferation [15].

We have previously used this mouse model of menstruation to investigate the role of uterine macrophages during endometrial repair which identified distinct monocyte and macrophage populations with spatially discrete distributions[16]. Specifically, immunohistochemistry analysis revealed that monocyte-derived macrophages were located in areas of the endometrium actively undergoing repair whilst tissue-resident macrophages were restricted to areas that have previously undergone repair [16]. Macrophages have been shown to contribute to endometrial shedding through the production of matrix metalloproteinases that degrade the extracellular matrix and through the production of vascular endothelial growth factor [17, 18]. Neutrophils are abundant during menstruation and are associated with apoptosis and cytokine production but whether they contribute to endometrial repair has not been explored [7, 8]. Cytokines and growth factors that support tissue breakdown and repair include pro-inflammatory cytokines (IL-1β, TNF-α, IL-6)[19, 20], chemokines (CCL2, CCL5, CXCL8) [8, 21] and regulators of matrix degradation and remodelling (MIF, MMPs, prostaglandins, VEGF) [1, 22, 23]. However, expression of these mediators has mostly been reported in whole tissues, limiting interpretation of cell-specific factors. Although immune cell types and mediators in menstruation have been profiled to some extent, deeper phenotyping of endometrial repair processes is needed to advance our understanding of menstrual physiology.

Menstruation is a complex and dynamic inflammatory process, but the contribution of distinct immune cell subsets and mediators remains poorly understood. To address this, we performed detailed phenotypic characterisation of uterine immune cells in a mouse model of simulated menstruation scRNAseq, flow cytometry and multiplex immunohistochemistry. We describe temporal and spatial association of myeloid cells with regions of tissue repair and highlight putative repair-associated signalling pathways associated with these cell types. Collectively, these data provide new insights into inflammatory regulation of menstruation and identify key pathways could be future therapeutic targets in menstrual disorders.

## Materials and Methods

### Mouse model of simulated menstruation

All animal work was carried out according to best practise in line with the Animal Welfare and Ethical Review Body (AWERB) and under licensed approval from the UK Home Office. The mouse model of simulated menstruation was conducted in accordance with standard protocols that have been described in detail in previous studies[8, 15, 16, 24, 25]. Briefly, female mice between eight to ten weeks of age underwent dorsal bilateral ovariectomy (d0) and were given daily injections of 17β-oestradiol (E2) in sesame oil (100ng/100µl, d7-9). A progesterone (P4)-secreting pellet was inserted subcutaneously on d13; mice also received daily injections of E2 (5ng/100µl; d13-15). On d15, decidualisation was induced by trans-cervical introduction of sesame oil into the uterine lumen (40µl, d15). Removal of the P4 pellet (d19) results in a rapid decline in progesterone, stimulating the breakdown and shedding of the decidual mass which simulates human menstruation in the mouse. Uterine tissues were collected at various time points following progesterone withdrawal (0-, 12-, 24-, and 48 hr) as well as from age-matched mice (∼3-month-old, oestrus cycling). A schematic of the mouse model of simulated menstruation and gross morphological changes that occur in the endometrium during phases of tissue breakdown, repair and remodelling can be seen in figure S1.

### Bioinformatics analysis of single cell RNA sequencing data

ScRNAseq analysis was performed on total leukocytes (CD45^+^ live singlets) isolated from uterine tissues 24-(n=4) and 48 hrs (n=4) following progesterone withdrawal, representing time points of endometrial tissue repair and remodelling respectively, and compared to age-matched cycling/control tissues (n=8). CD45^+^ cells were recovered using fluorescence-activated cell sorting (FACS), pooling 25,000 cells per sample to give a total of 100,000 cells for input to the 24/48hr dataset and 200,000 cells for input to the cycling dataset. Isolated cell suspensions were counted, and viability confirmed to be >85% using a TC20 Automated Cell Counter (BioRad, Cat. No. 1450102). Cell suspensions were processed using the 10x Chromium Controller (10x Genomics, USA) to partition single cells into Gel bead-in-Emulsions (GEMs) containing unique 10x barcodes and cDNA libraries generated following standard protocol

ls as previously described[24, 25]. Sequencing was carried out on the Illumina NovaSeq platform using bespoke 10x parameters, according to standard protocols (Edinburgh Genomics: http://genomics.ed.ac.uk/, Edinburgh).

Raw data files were processed using 10x Cell Ranger (version 2.0.1, https://www.10xgenomics.com) and downstream QC, clustering and gene expression analysis performed in *Seurat [26]* (Version 3) following analysis pipelines described previously[24, 25]. Data were filtered based on standard QC metrics: number of unique genes detected in each cell (nFeature_RNA >200 & <5000); the percentage of mitochondrial genes (<10%); and the percentage ribosomal genes (<10%). The Scrublet python package was used to further identify and remove putative doublets from scRNAseq data[27]. Individual datasets were combined using Seurat’s ‘anchor-based’ integration workflow to facilitate accurate comparative analysis between experimental conditions. Following integration, data were normalised and scaled prior to unsupervised clustering based on PCA analysis of the most variably expressed genes using a graph-based approach (‘FindNeighbours’, ‘FindClusters). A quantitative method was used alongside the ‘ElbowPlot’ function to determine the number of principle components required to capture the majority of the variance in the data[28]. Resultant clusters were visualised using the uniform manifold approximation and projection (UMAP) method and differential gene expression analysis was performed (‘FindAllMarkers’) to identify cluster specific gene signatures using the non-parametric Wilcoxon rank sum test and p-value threshold of <0.05. Over-represented functional annotations associated with the differentially expressed genes (DEGs) were detected using the clusterProfiler package[29] using core functions to interpret data in the context of biological function, pathways, and networks.

The R package CellChat [30] (version 2) was utilised to investigate cellular interactions between immune cell types. CellChat estimates the probability of cell-cell communication via the integration of single cell gene expression data with a database (CellChatDB) of known signalling ligands and receptors generated from the literature.

The data discussed in this study will be deposited in NCBI’s Gene Expression Omnibus on publication [31].

### Flow cytometry

Uterine tissue samples were dissected and transferred to PBS kept at 4°C prior to the tissue digestion protocol. Individual uterine horns were analysed as distinct samples given the potential for each to exhibit varying degrees of decidualisation in response to the stimulus. Tissue processing for flow cytometry analysis was performed as described previously [24, 25]. Briefly, uterine tissues were manually dissociated and incubated with collagenase (1mg/ml) and DNase (0.1mg/ml) for 30 min at 37°C. Tissues were further dispersed and washed in FACS buffer (PBS Ca2-Mg2-: 5% charcoal stripped foetal calf serum (CSFCS – Cat. No. 12676029), 2mM EDTA) and subsequently strained through 70μm and 40μm cell strainers. Cell suspensions were centrifuged at 400rcf for 5 min and cell pellets re-suspended in 1ml ACK lysing buffer (Gibco, Cat. No. A10492-01) for 1 min then washed. Uterine cell suspensions were incubated with LIVE/DEAD Red (Life Technologies Ltd, L34971) for 30 min on ice followed by incubation with Mouse TruStain FcXTM (Biolegend 101319; 1:1000) for 10 min on ice. Cell suspensions were subsequently incubated with optimised dilutions of fluorescently conjugated extracellular antibodies for 30 min on ice, as detailed in Table 1 –- Figure 2.

Following antibody incubation, single cell suspensions were washed and re-suspended in FACS buffer at 4°C and analysed using a Cytek Aurora (Cytek Biosciences) and SpectroFlo software. Data analysis was performed using FlowJo analysis software (FlowJo LLC, Oregon, USA).

### Tissue recovery, processing and Haematoxylin and Eosin staining

Mouse uteri were fixed in 10% neutral buffered formalin for 24 hr at 4°C before transfer into 70% ethanol, at which point they could be stored long term. Tissues were then processed according to standard protocols for paraffin embedding by the University of Edinburgh Histology Facility. Sections from formalin-fixed paraffin-embedded (FFPE) blocks were cut at 5um thickness using a microtome and the sections were placed on Xtra adhesive pre-cleaned micro slides (Surgipath, Leica Biosystems) and placed in at 60°C for 15 min prior to staining. For visualisation of tissue morphology, staining was performed with haematoxylin and eosin.

### Immunohistochemistry and Immunofluorescence

For immunohistochemistry and immunofluorescence, FFPE uterine sections of 5um thickness were deparaffinised and rehydrated using a series of xylene and alcohols followed by antigen retrieval in Citrate retrieval buffer (pH6) for 5 min in an Instant Pot. Following washes with tap water, the sections were incubated with 3% (v/v) hydrogen peroxide solution in PBS for 30 min at room temperature, followed by several blocking steps with serum from the species the secondary antibody was raised in, to avoid nonspecific binding of antibodies. Sections were incubated with the primary antibody diluted in serum blocking solution overnight at 4°C. Negative controls were included, in which the primary antibody was omitted to identify any nonspecific binding by the secondary antibody. Primary antibodies against CD3 (1:250 dilution; NB600-1441, Novus Biologicals), B220 (1:250 dilution; 14-0452-82, Invitrogen), CD14 (1:6000 dilution; ab221678, Abcam), CD64 (1:6000 dilution; 50086-R008, Sino Biological), Ly6G (1:10000 dilution; 127649, Biolegend) and DBA-lectin (1:1000 dilution; B-1035, Vector) were used in this study.

The following day, sections were washed with 0.05% Tween in PBS followed by an incubation with an HRP-conjugated secondary antibody (1:500 dilution) in serum blocking solution for 30 min at room temperature followed by an incubation with either DAB substrate for 5 min at room temperature (ab64238, Abcam) or Opal Polaris solution (FP1487001KT [Opal 520], FP1488001KT [Opal 570], FP1496001KT [Opal 650], Akoya Biosciences) for 10 min at room temperature. After staining, sections were washed and counterstained either with Haematoxylin or DAPI (1:3000 dilution; 10236276001, Sigma-Aldrich). For chromogenic immunohistochemistry, the sections were dehydrated in a series of alcohols and mounted on coverslips with Pertex mounting medium (SEA-0100-00A, CellPath), while for immunofluorescence, the sections were mounted on coverslips with Fluoromount (0100-01, Cambridge Bioscience). Slides were imaged with a slide scanner (Axioscan 7, Zeiss) or an inverted microscope (Axio Observer, Zeiss) and images were exported using the Zen 3.9 software (Zeiss).

### Statistical analysis

Statistical analysis was performed using Graphpad prism software (version 10). Brown-Forsythe and Welch tests were used to determine significance between experimental conditions in data that were normally distributed. Non-parametric testing was utilised where sample sizes were insufficient to confirm normality of data distribution or where the data was not normally distributed; Kruskal-Wallis tests were used to assess differences between experimental conditions. All data are presented as mean ± SD and criteria for significance is p<0.05.

## Results

### Single-cell RNA sequencing identifies distinct immune cell populations in the mouse uterus associated with endometrial tissue repair and remodelling

To investigate the phenotype and function of immune cells during endometrial repair, single cell RNA sequencing (scRNAseq) analysis was performed on CD45^+^ immune cells isolated from uterine tissues recovered at key time points of an established mouse model of menstruation: 24- and 48 hr post progesterone withdrawal and in age-matched controls (‘cycling’) (figure S1). These timepoints are representative of active tissue repair (24 hr, ‘repair’) and tissue remodelling (48 hr, ‘remodelling’). Following quality control and filtering of sequencing data, 13,498 CD45^+^ immune cells were detected across 3 experimental conditions (cycling: 5,379 cells; repair: 3,761 cells; and remodelling: 4,358 cells) and integrated for downstream analysis. Unsupervised clustering resulted in the identification of 8 transcriptionally distinct clusters of immune cells in uterine tissues: neutrophils, eosinophils, monocytes, macrophages, dendritic cells, NK cells, T cells and B cells (Figure 1A). Cell clusters were annotated using both the expression of established immune cell gene markers (Figure 1B) and analysis of the top 10 most differentially expressed genes (DEGs) per cluster (Figure 1C & 1D).

**Figure 1:**
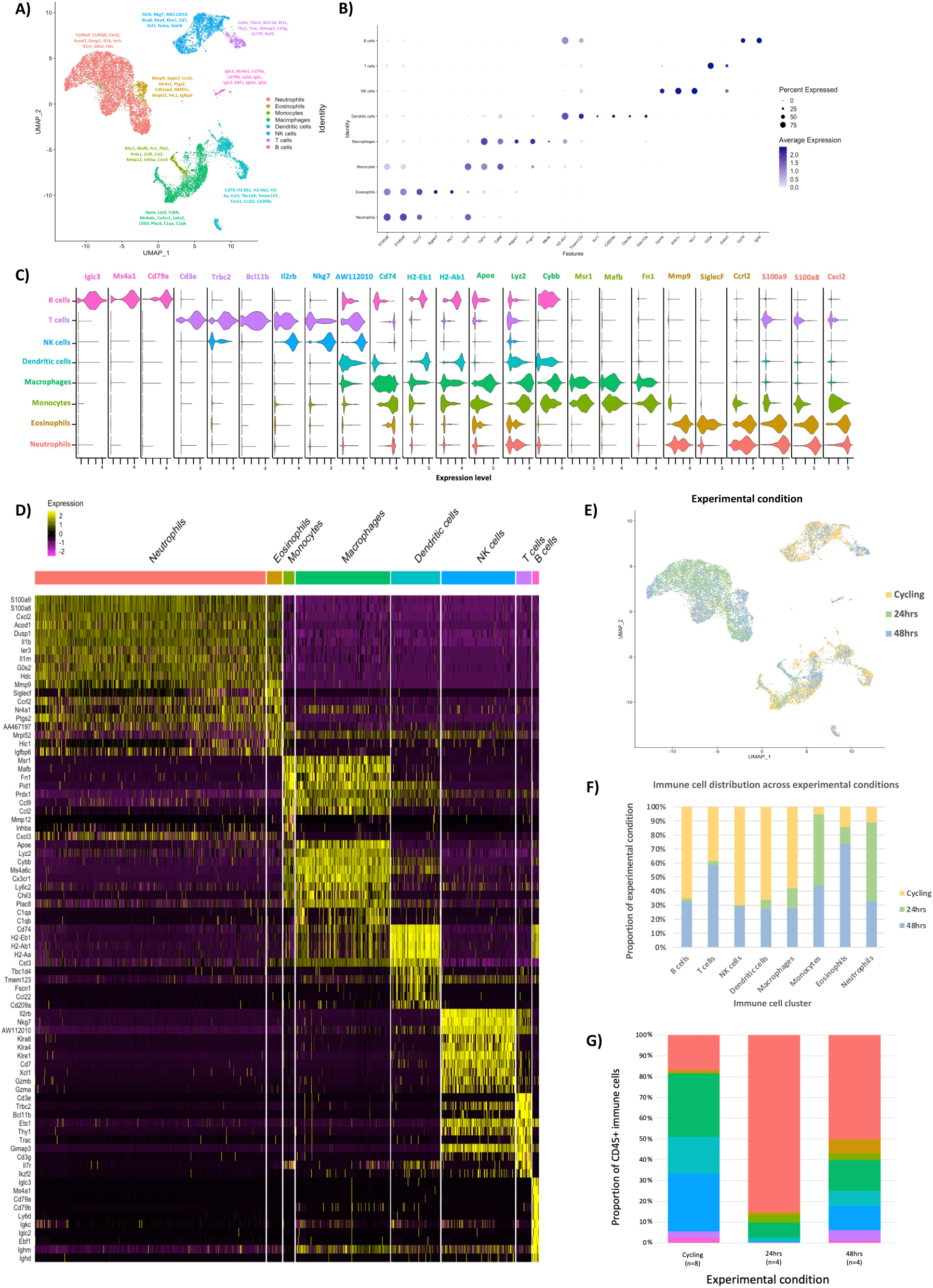
Single cell RNA sequencing analysis identifies 8 uterine immune cell populations in a mouse model of endometrial repair. **(A)** Uniform manifold approximation and projection (UMAP) visualisation: population restricted CD45^+^ immune cells isolated from mouse uterine tissue samples taken from steady state control mice (cycling, n=8) and menstruating mice 24-(endometrial repair, n=4) and 48 hr (endometrial remodelling, n=4) after progesterone withdrawal. **(B)** Dot plot: Expression of canonical immune cell markers: neutrophils (*S100a8*, *S100a9*, *Cxcr2*), eosinophils (*SiglecF*, *Hic1*), monocytes (*Cd14*), macrophages (*Csfr1, Cd68, Adgre1*, *Fcgr1, Mertk*), dendritic cells (*H2-Ab1, Tmem123, Xcr1, Cd209a, Clec9a, Clec10a*), natural killer cells (*Gzmb, Klrb1c, Ncr1*), T cells (*Cd3e, Gata3*), and B cells (*Cd19, Ighd*). Dot size: percentage of cells in each cluster expressing the gene; Dot colour: average expression per cluster from low (lavender) to high (dark blue). **(C)** Violin plot: Expression of the top 3 differentially expressed genes in each cell cluster displayed as log (TPM^+^1). TPM= transcript count per million. **(D)** Heatmap (yellow, high; purple, low): Top 10 differentially expressed genes in each cell cluster (logFC > 0.5, p-value <0.05, Wilcoxon rank-sum test). Top axis is colour coded and named by immune cell type. **(E)** Feature plot: Distribution of cells from each experimental condition. **(F)** Bar plot: Summary of immune cell origin across experimental conditions. Each colour represents each experimental condition, and the y-axis denotes the contribution of each experimental condition to each immune cell cluster. **(G)** Stacked bar plot: Single cell RNA sequencing analysis of the relative abundance of uterine immune cell populations during endometrial repair. Each colour represents a distinct immune cell type, and the y-axis denotes the percentage of total CD45^+^ immune cells.

Analysis of immune cell clusters relative to experimental condition identified distinct changes in immune cell composition during endometrial repair and remodelling. Innate immune populations including neutrophils, monocytes and to a lesser degree macrophages, dendritic cells and eosinophils were predominant in the ‘repair’ dataset, while adaptive immune populations such as T, B and NK cells, were largely only present in the ‘cycling’ and ‘remodelling’ data (Figure 1E & 1F). Relative abundance analysis within each experimental condition showed changes in uterine immune cell composition in response to endometrial repair (Figure 1G). Most notable was an increase in the proportion of neutrophils and monocytes during active endometrial repair (24-hour) which remained elevated during the remodelling phase (48-hour).

Unbiased gene ontology (GO) analysis was used to identify biological processes associated with the transcriptome of each immune cell cluster in the combined dataset. Top processes associated with subsets were consistent with their phenotypic annotation, further validating their identification as transcriptionally and functionally discrete immune cells (Figure S2A). Monocytes were associated with ‘wound healing’ (Figure S2A(v)) macrophages with phagocytosis (Figure S2A(vi)) and neutrophils with cell chemotaxis/migration (Figure S2A(vii)).

To explore the functional role of monocytes, macrophages and neutrophils in endometrial repair and remodelling, the expression of gene sets associated with essential tissue repair processes, generated from literature searches and MsigDB [32], were interrogated in scRNAseq data (Figure S2B). Neutrophils were found to have the highest expression of genes associated with NETosis, chemoattraction and certain remodelling enzymes while macrophages and monocytes expressed genes associated with phagocytosis. Notably, when assessing the expression of genes associated with ‘wound healing’ each cell type showed expression of a distinct subset of genes (monocytes: *Anxa5*, *Cav1*, *F10*, *Fn1*, *Pdpn*, *Sdc1*, *Sdc4, Thbd*; macrophages: *Apoe*, *Axl*, *C3*, *Ccr2*, *Gnas*; neutrophils: *Slc11a1, Cd9, Lrg1, Plaur, S100a9*) (Figure S3B). Taken together, GO analyses suggest that myeloid cells are predominant during endometrial tissue repair and may have key role in wound healing following menstruation.

### Flow cytometry analysis confirms dynamic changes in immune cell composition and the predominance of myeloid cells during endometrial repair

To validate putative repair-associated changes in uterine immune cell composition identified by *in silico* analyses, flow cytometry was performed on uterine tissues collected over a comprehensive time course of endometrial repair using the mouse model of menstruation (figure S1). A flow cytometry panel was designed to interrogate the immune cell populations identified by scRNAseq using phenotypic markers as outlined in Supplementary Table 1.

Using the gating strategy outlined in figure 2A, flow cytometry analysis corroborated the immune cell composition identified by scRNAseq analysis in cycling, repair (24 hr) and remodelling (48 hr) tissues. This was complemented by analysis of additional time points which provided resolution on immune cell dynamics prior to and during tissue breakdown (Figure 2B & 2C). At 0 hr (prior to breakdown), there was an increase in the proportion of monocytes within the uterus and a decrease in the proportion of NK cells and dendritic cells (DCs) when compared to cycling control. At 12 hr during tissue breakdown there was a large compositional shift characterised by a significant increase in the proportion of neutrophils in the uterus, accounting for 87.82% ± 14.06 of all CD45^+^ immune cells at this timepoint. The proportion of neutrophils peaked at 24 hr (91.86% ± 2.60 of CD45^+^ cells) during repair and neutrophils were also the most abundant cell type at 48 hr (55.7% ± 12.72 of CD45^+^ cells), with the numbers of T cells, NK cells and macrophages also increasing at 48 hr (Figure 2B & 2C).

**Figure 2:**
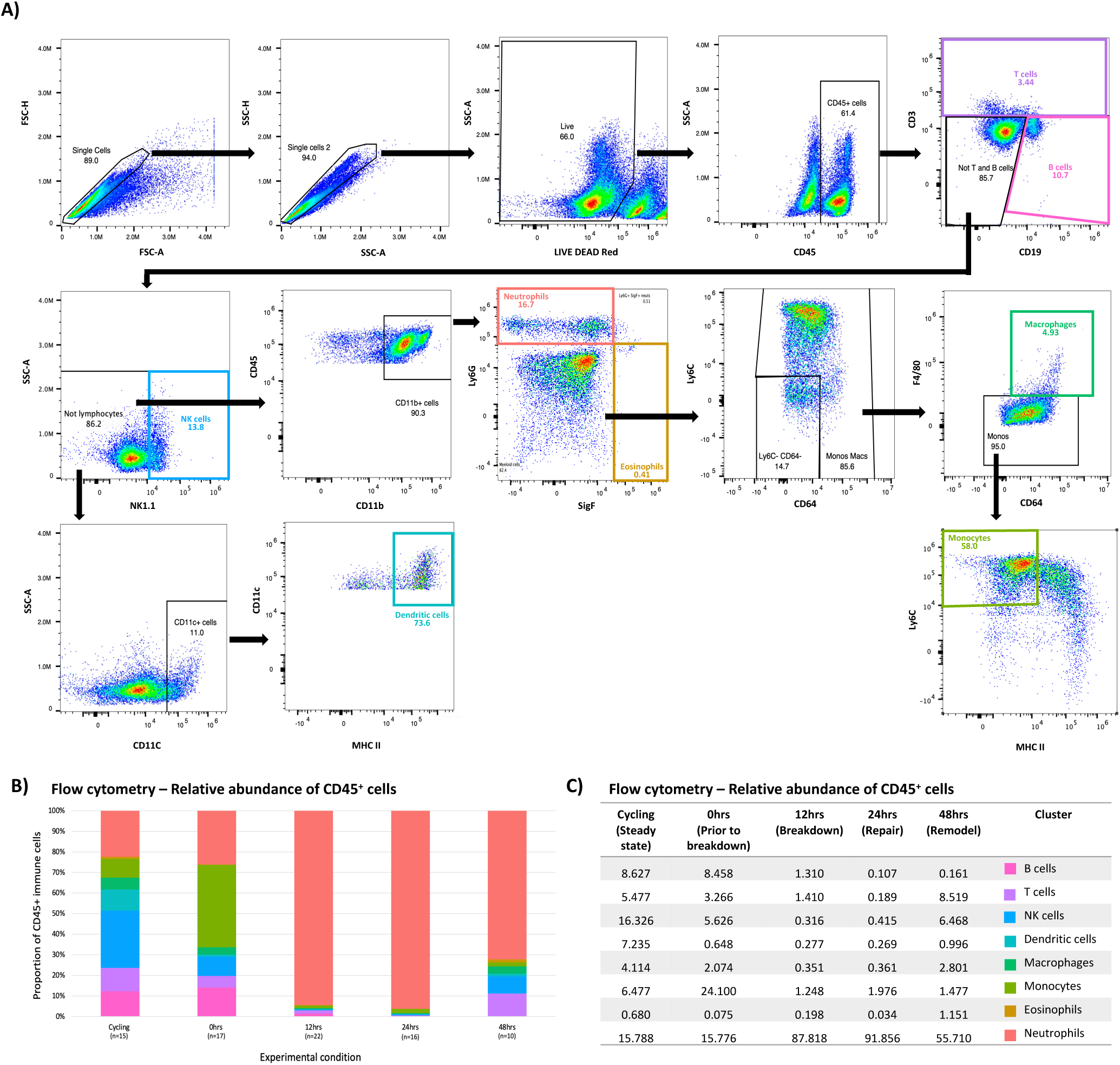
Flow cytometry analysis reveals dynamics changes in uterine immune cell composition during a mouse model of endometrial repair and remodelling. **(A)** Flow cytometry gating strategy employed to identify and quantify distinct immune cell populations in uterine tissue digests. **(B)** Stacked bar plot: relative abundance of uterine immune cell populations across phases of endometrial repair and remodelling (0-,12-, 24-, 48 hr following progesterone withdrawal) compared to cycling/steady state control tissue: colours represent distinct immune cell types, and the y-axis denotes the percentage of total CD45^+^ immune cells in each dataset. **(C)** Table: Quantification of flow cytometry analysis of the relative abundance of uterine immune cell populations across phases of endometrial repair and remodelling: 0 hr (n=17), 12 hr (n=22), 24 hr (n=16) and 48 hr (n=10) following progesterone withdrawal compared to cycling/steady state control tissue (n=15).

### Lymphocyte populations are infrequent during endometrial repair

T cells (CD45^+^ Ly6G^−^ NK1.1^−^ CD19^−^ CD3^+^) were quantified in uterine tissue digests using flow cytometry (Figure 3A). Both the relative abundance (percentage of CD45^+^ cells) and absolute numbers of T cells were significantly increased during tissue remodelling (48 hr; 8.52% ± 4.71 and 47.87 count/mg ± 27.80) when compared to earlier phases of breakdown (12 hr; 1.72% ± 3.84 and 8.06 count/mg ± 9.67) and repair (24 hr; 0.19% ± 0.13 and 11.77 count/mg ± 13.23) (Figure 3B & 3C). IHC was performed on tissue samples from matching time points/conditions. CD3^+^ T cells were detected in the endometrium and myometrium of both 24- and 48-hour tissues (Figure 3D; black arrows & Figure S4A). T cells were more readily detected in 48-hour tissues but there was no spatially distinct distribution associated with different time-points (Figure 3D; black arrows & Figure S4A).

**Figure 3:**
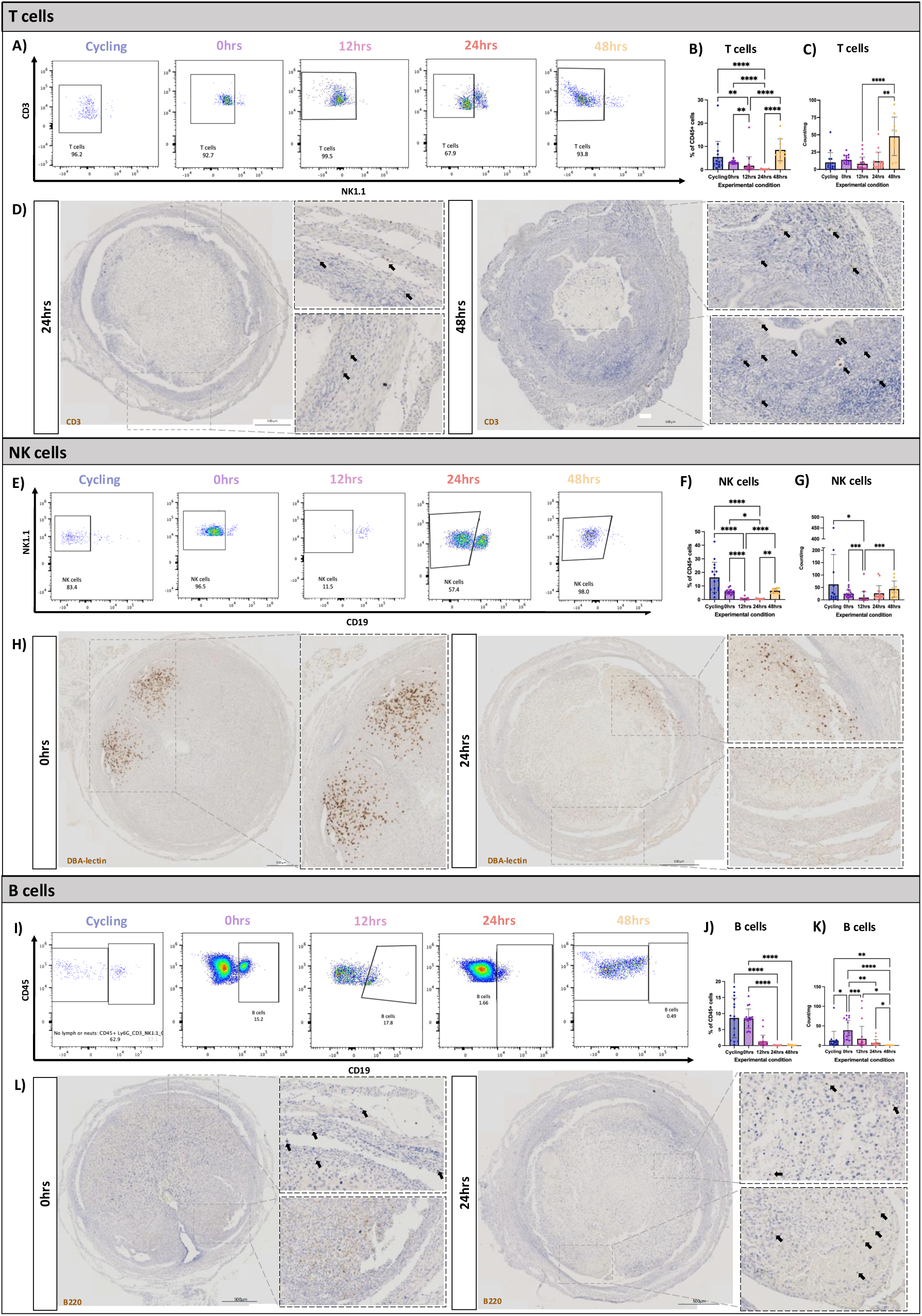
Flow cytometry and immunohistochemistry reveal dynamic changes in uterine lymphocyte populations during endometrial repair using a mouse model of simulated menstruation. **(A)** Population restricted FC analysis to identify CD45^+^ Ly6G^−^NK1.1^−^ CD19^−^ CD3^+^ T cells in uterine tissue digests. **(B)** Bar plot, FC quantification: Relative abundance of T cells calculated as a percentage of total CD45^+^ cells per uterine horn. **(C)** Bar plot, FC quantification: Absolute counts of T cells per uterine horn. **(D)** Immunohistochemistry: detection of CD3^+^ T cells (black arrows) in uterine tissues 24- and 48 hr after progesterone withdrawal (representative images: 24 hr n=5, 48 hr n=6, scale bar= 50µm). **(E)** Population restricted FC analysis to identify CD45^+^ Ly6G^−^ CD3^−^ CD19^−^ NK1.1^+^ NK cells in uterine tissue digests. **(F)** Bar plot, FC quantification: Relative abundance of NK cells calculated as a percentage of total CD45^+^ cells per uterine horn. **(G)** Bar plot, FC quantification: Absolute counts of NK cells per uterine horn. **(H)** Immunohistochemistry: detection of DBA-bound NK cells in uterine tissues 24- and 48 hr after progesterone withdrawal (representative images: 24 hr n=4, 48 hr n=4, scale bar= 50µm). **(I)** Population restricted FC analysis to identify CD45^+^ Ly6G^−^ CD3^−^ NK1.1^−^ CD19^+^ B cells in uterine tissue digests **(J)** Bar plot, FC quantification: Relative abundance of B cells calculated as a percentage of total CD45^+^ cells per uterine horn. **(K)** Bar plot, FC quantification: Absolute counts of B cells per uterine horn. **(L)** Immunohistochemistry: detection of B220^+^ B cells (black arrows) in uterine tissues 24- and 48 hr after progesterone withdrawal (representative images: 24 hr n=4, 48 hr n=4, scale bar= 50µm). (FC analysis groups: control (n=15), 0 hr (prior to breakdown, n=17), 12 hr (tissue breakdown, n=22), 24 hr (tissue repair, n=16) and 48 hr (tissue remodelling, n=10) after progesterone withdrawal; statistical comparisons were made using Kruskal-Wallis tests with multiple comparisons. * p< 0.05, ** p < 0.01, *** p < 0.001, **** p < 0.0001).

NK cells (CD45^+^ Ly6G^−^ CD3^−^ CD19^−^ NK1.1^+^) were more readily detected (Figure 3E) and abundant in cycling mice (62.35 count/mg ± 120.30), numbers were lowest during breakdown (12 hr; 9.069 count/mg ± 23.66) and repair (24 hr; 26.44 count/mg ± 30.64) but more abundant in remodelling tissues (48 hr; 43.35 count/mg ± 32.52) (Figure 3G). Detection of NK cells using DBA-lectin in uterine tissue sections located NK cells to the decidualised tissue prior to breakdown (0-hour), in close proximity to the luminal epithelium and adjacent to the decidual-endometrial boundary. In 24-hour tissues NK cells were infrequent, located predominantly in the shed decidual tissue (Figure 3H) and in 48-hour tissues very few NK cells could be detected (Figure S4B).

Quantitative analysis of B cells (CD45^+^ Ly6G^−^ CD3^−^ NK1.1^−^ CD19^+^) demonstrated that B cells are abundant in the uterus prior to breakdown (0 hr; 38.66 count/mg ± 24.97) but significantly decreased thereafter (Figure 3J & 3K). Detection of B220^+^ B cells in uterine tissue sections by IHC showed B cells located in both the endometrium and myometrium prior to breakdown (Figure 3L; 0 hr, black arrows) but they were infrequent during repair and remodelling (Figure 3L; black arrows & Figure S4C).

### Myeloid cell populations are dynamically regulated during endometrial breakdown, repair, and remodelling

Flow cytometry and immunohistochemistry analyses demonstrated changes in the abundance and location of key myeloid cell types in the uterus during endometrial repair. Monocytes (Ly6C^+^ MHCII^−^; Figure 4A) were the most abundant immune cell present in the uterus prior to breakdown (0-hour; 103.8 ± 50.46) (Figure 4B & 4C). The number of monocytes in the uterus was then significantly decreased during breakdown when compared to prior to breakdown (0 hr; 103.80 count/mg ± 50.46 vs 12 hr; 25.40 count/mg ± 54.57, p < 0.0001). Monocytes were significantly increased during repair (12 hr; 25.40 count/mg ± 54.57 vs 24 hr; 101.1 count/mg ± 49.67, p < 0.0001) and significantly decreased during remodelling (24 hr; 101.1 count/mg ± 49.67 vs 48 hr; 10.44 count/mg ± 8.27, p = 0.0018) (Figure 4C). Immunofluorescent staining was performed using an anti-CD14 antibody to investigate spatial distribution of monocytes in uterine tissues. CD14^+^ cells were predominantly localised to the boundary between the underlying endometrium and the detached decidual tissue during breakdown (12 hr; Figure S6A) and subsequent repair (24 hr; Figure 4J and Figure S6A).

**Figure 4:**
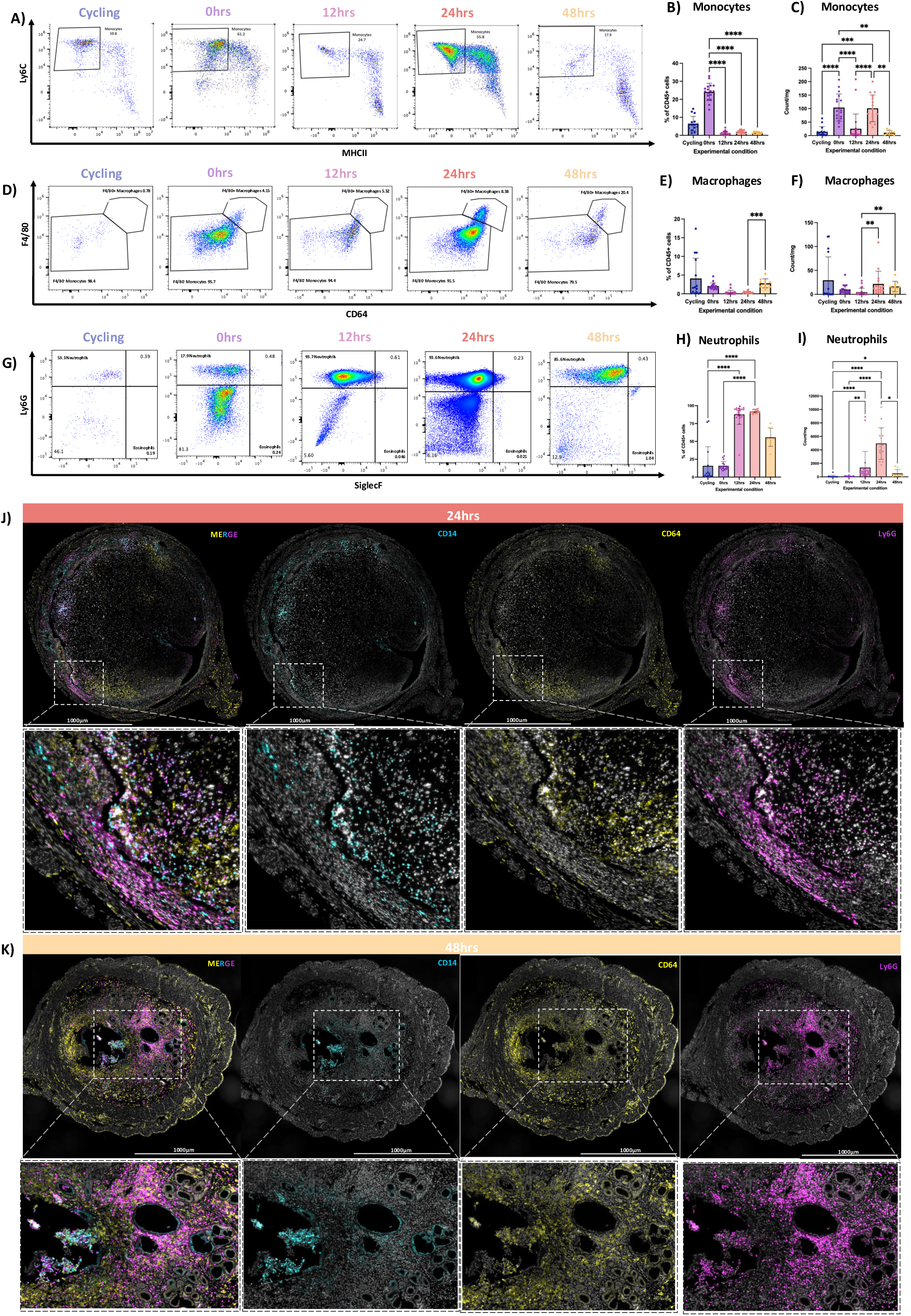
Flow cytometry and immunohistochemistry reveal dynamic changes in uterine myeloid cell populations during endometrial repair using a mouse model of simulated menstruation. **(A)** Population restricted FC analysis to identify CD45^+^ CD3^−^ CD19^−^NK1.1^−^ CD11b^+^ Ly6G^−^ SiglecF^−^ CD64^−^ Ly6C^+^ MHCII^−^ monocytes in uterine tissue digests. **(B)** Bar plot, FC quantification: Relative abundance of monocytes calculated as a percentage of total CD45^+^ cells per uterine horn. **(C)** Bar plot, FC quantification: Absolute counts of monocytes per uterine horn. **(D)** Population restricted FC analysis of CD45^+^ CD3^−^ CD19^−^NK1.1^−^ CD11b^+^ Ly6G^−^ SiglecF^−^ CD64^+^ Ly6C^−^ F4/80^+^ MHCII^+^ macrophages in uterine tissue digests. **(E)** Bar plot, FC quantification: Relative abundance of macrophages calculated as a percentage of total CD45^+^ cells per uterine horn. **(F)** Bar plot, FC quantification: Absolute counts of macrophages per uterine horn. **(G)** Population restricted FC analysis of CD45^+^ CD3^−^CD19^−^ NK1.1^−^ CD11b^+^ Ly6G^+^ SiglecF^−^ neutrophils in uterine tissue digests. **(H)** Bar plot, FC quantification: Relative abundance of neutrophils calculated as a percentage of total CD45^+^ cells per uterine horn. **(I)** Bar plot, FC quantification: Absolute counts of neutrophils per uterine horn. (FC analysis groups: control (n=15), 0 hr (prior to breakdown, n=17), 12 hr (tissue breakdown, n=22), 24 hr (tissue repair, n=16) and 48 hr (tissue remodelling, n=10) after progesterone withdrawal; statistical comparisons were made using Kruskal-Wallis tests with multiple comparisons. * p< 0.05, ** p < 0.01, *** p < 0.001, **** p < 0.0001). **(J-K)** Immunohistochemical detection of CD14^+^ monocytes (cyan), CD64^+^ macrophages (yellow) and Ly6G^+^ neutrophils (magenta) (merged and split channel) in uterine tissue cross sections at **(J)** 24 hr (repair, n=5) and **(K)** 48 hr (remodelling, n=3) following progesterone withdrawal (merged and split channel representative images, scale bar = 1000µm).

Macrophages counts were significantly decreased during tissue breakdown when compared to tissue repair (12 hr; 4.71 count/mg ± 8.17 vs 24 hr; 21.78 count/mg ± 26.47, p = 0.0013) and remodelling time points (12 hr; 4.71 count/mg ± 8.17 vs 48 hr; 16.61 count/mg ± 10.69, p < 0.0001) (Figure 4F). Macrophages (F4/80^+^CD64^+^; Figure 4D) were decreased during endometrial repair but increased in the remodelling timepoint (24 hr; 0.36% ± 0.28 vs 48 hr; 2.80% ± 1.22, p = 0.004) (Figure 4E). CD64^+^ macrophages were detected by immunohistochemistry in all uterine tissue compartments throughout breakdown and repair (0-24 hr; Figure S6B), in both the decidualised and non-decidualised endometrium, as well as the myometrium. In 48-hour tissues, macrophages were most abundant in the stroma of the remodelled endometrium (Figure 4K and Figure S6B).

Neutrophils (Ly6G^+^SiglecF^−^, Figure 4G) were the most abundant immune cell type present in the uterus during breakdown (12 hr; 87.82% ± 14.06), repair (24 hr; 91.86% ± 2.60), and remodelling (48 hr; 55.71% ± 12.72) (Figure 4H). Neutrophils counts in the uterus were significantly increased during breakdown compared to cycling controls (12 hr; 1374 count/mg ± 2398 vs cycling; 56.25 count/mg ± 175.80, p < 0.0001). Neutrophil abundance peaked during endometrial repair (24 hr; 4946 count/mg ± 2344) and although neutrophil numbers were relatively decreased during remodelling they were still significantly increased compared to cycling controls (Figure 4I; cycling; 56.25 count/mg ± 175.80 vs 48 hr; 504.5 count/mg ± 570.6, p = 0.016). Immunofluorescent staining identified abundant Ly6G^+^ neutrophils in uterine tissues during breakdown (12 hr) and repair (24 hr) specifically localised to the boundary between the detached decidual tissue and the underlying endometrium (Figure S6C). In 48-hour tissues, Ly6G^+^ neutrophils were readily detected throughout the remodelled endometrium (Figure S6C).

Dendritic cells (DCs; CD11c^+^MHCII^+^; Figure S5A) were infrequent during tissue breakdown (12 hr; 5.63 count/mg ± 11.91) but were modestly increased during repair (24 hr; 15.80 count/mg ± 13.51) and remodelling (48 hr; 6.59 count/mg ± 6.019) (Figure S5B & S5C). Eosinophils (Ly6G^−^ SiglecF^+^; Figure S5D) were infrequent in cycling tissues (3.26 count/mg ± 6.22) and during phases of tissue breakdown (12 hr; 2.14 count/mg ± 5.08) and repair (24 hr; 2.43 count/mg ± 4.20). Notably, eosinophils were significantly increased during remodelling (48 hr; 17.60 count/mg ± 20.43) but were infrequent overall (Figure S5F). Due to their low abundance, DCs and eosinophils were not analysed by IHC.

### Monocytes, neutrophils, and macrophages co-localise to areas of breakdown and tissue repair

To understand the distribution of key myeloid cell types relative to each other in uterine tissue sections, multiplex IHC for the detection of CD14 (monocytes), CD64 (macrophages) and Ly6G (neutrophils) was performed using uterine tissues collected from across the time course (Figure 4J-K, Figure S6D). Prior to tissue breakdown (0 hr), relatively few CD14^+^ or Ly6G^+^ cells could be detected but CD64^+^ macrophages were abundant throughout the tissue (Figure S6D). At 12 hr, during tissue breakdown, CD14^+^ monocytes and Ly6G^+^ neutrophils were detected clustered within the same regions of the tissue and were localised to regions of shedding, where decidualised tissue was detaching, as well as in the underlying residual endometrium (Figure S5D). CD64^+^ macrophages were located predominantly in the outer myometrial layers at 12 hr and only diffusely detected within regions of tissue breakdown (Figure S6D).

At 24 hr, CD14^+^ monocytes and Ly6G^+^ neutrophils were identified in the detaching decidual tissue but were also detected in regions of underlying endometrium undergoing tissue repair (Figure 4J & S6D). CD64^+^ macrophages were also detected throughout the repairing endometrium as well as in the decidual tissue, in association with regions where detachment had not yet taken place (Figure 4J & S6D). At this time point, monocytes, neutrophils and macrophages were detected within regions of active tissue repair but were less frequent in areas of intact tissue. At 48 hr the decidual tissue is almost completely expelled, the luminal epithelium is largely restored, and the underlying stroma is undergoing remodelling. CD14^+^ monocytes were less frequent at 48 hr and localised only to regions of decidual detachment on the endometrial surface, while Ly6G^+^ neutrophils were abundant and dispersed throughout the remodelling endometrium (Figure 4K & S6D). Notably, CD64^+^ macrophages were more abundant at 48 hr with an apparent expansion throughout the remodelling endometrium in regions both proximal and distal to sites of recent tissue repair (Figure 4K & S6D).

### CellChat analysis of scRNAseq data identifies monocyte, macrophage and neutrophil signalling pathways associated with endometrial repair

Given the association of monocytes, macrophages and neutrophils with tissue repair and remodelling in the endometrium, further *in silico* analyses were performed to infer putative signalling pathways and interactions associated with these cell types. CellChat analysis of ligand-receptor expression was performed on scRNAseq data, and immune cell interactions were assessed. Monocytes, macrophages and neutrophils were identified as both dominant senders (high ligand expression) and dominant receivers (high receptor expression) in the communication network, scoring highly in both outgoing and incoming interaction strength (Figure 5A). The top signalling pathways found to contribute to outgoing/incoming signalling across all immune cell types were CCL, CXCL, MIF, FN1, SPP1 and THBS (Figure 5B). Analysis of communication patterns between cell types for these signalling pathways highlighted the predominance of monocytes (CCL, CXCL, MIF, FN1, SPP1 and THBS), macrophages (CCL, MIF and FN1) and neutrophils (CCL, CXCL, and THBS) as key contributors to cell communication of repair-associated signalling pathways (Figure 5C).

**Figure 5.**
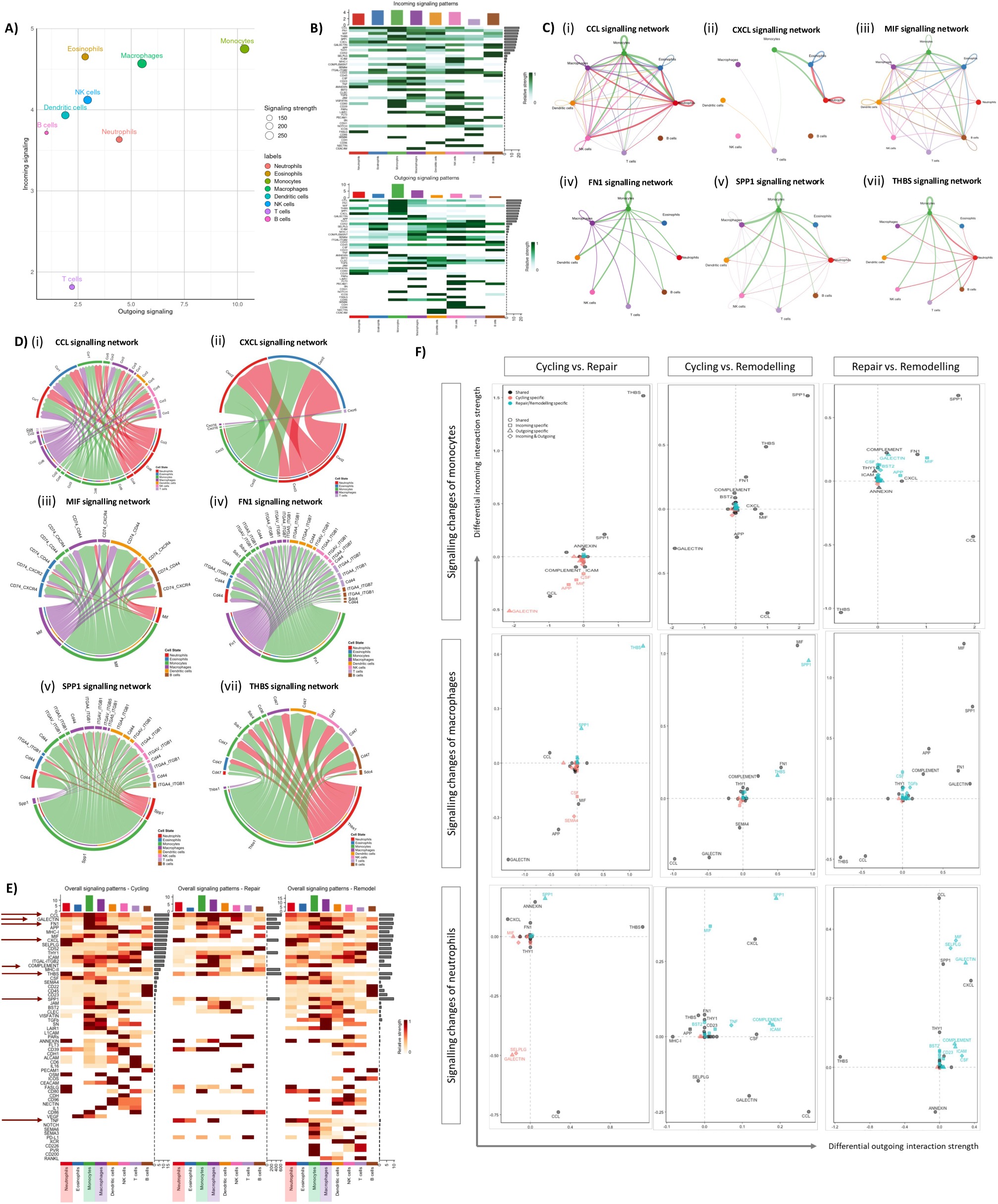
CellChat analysis of scRNAseq immune cell data infers intercellular communication networks associated with endometrial repair and remodelling. **(A)** Scatterplot: visualisation of overall signalling role of each cell group as determined by total outgoing and incoming communication probability (dot size = total number of inferred links, dot colour = different cell groups). **(B)** Heatmap: Outgoing and incoming signalling patterns relative to cell type identifying top pathways in scRANseq data. **(C)** Circle plot: inter- and intracellular communication networks between immune cell types for **(i)** CCL, **(ii)** CXCL, **(iii)** MIF, **(iv)** FN1, **(v)** SPP1, **(vii)** THBS signalling pathways (circle size = number of cells in group, connection width = probability of interaction). **(D)** Chord diagram: ligand-receptor pairs and their weighted interactions between immune cells for **(i)** CCL, **(ii)** CXCL, **(iii)** MIF, **(iv)** FN1, **(v)** SPP1, **(vii)** THBS signalling pathways (chord width = strength of interactions). **(E)** Heatmap: Overall signalling patterns of immune cells relative to experimental condition, top 8 signalling pathways highlighted with arrows (intensity of colour= relative signalling strength). **(F)** Scatterplot: pairwise comparison of relative strength of signalling pathways in monocytes, macrophages and neutrophils between experimental data set: cycling vs repair; cycling vs remodelling; repair vs remodelling.

The relationship between specific ligand-receptor pairs and immune cell subsets is summarised in the chord diagrams for each signalling pathway, where the width of each chord corresponds to the strength of the interaction (Figure 5D). The relationship between different CCL chemokine ligands and corresponding receptors showed a predominance of ligand expression associated with monocytes (*Ccl2*, *Ccl3*, *Ccl4*, *Ccl6* and *Ccl9*), macrophages (*Ccl2*, *Ccl3*, *Ccl4*, *Ccl6 and Ccl9*) and neutrophils (*Ccl3*, *Ccl6* and *Ccl9*) (Figure 5Di). Crosstalk in CXCL pathway was predominantly from monocytes and neutrophils signalling *Cxcl2* and *Cxcl3* to *Cxcr2* on neutrophils and eosinophils (Figure 5Dii). *Mif* and *Fn1* signalled from monocytes and macrophages to corresponding receptors on multiple cell types (Figure 5Diii and iv). *Spp1* ligand was predominantly associated with monocytes signalling to distinct reciprocal receptors including *Cd44* on neutrophils and integrins on monocytes and macrophages(Figure 5Dv). Similarly, *Thbs1* signalling was predominantly from monocytes and neutrophils which signalled via *Cd47* on all immune cell types in the dataset (Figure 5Dv and vi).

To examine the association of these signalling pathways more discretely, the top overall signalling pathways were assessed for each experimental condition (Figure 5E). While similar signalling pathways ranked highly, their association with specific immune cell subsets shifted according to experimental conditions. The relative strength of THBS signalling was strongest for neutrophils in cycling, but more strongly associated with monocytes in repair and remodelling tissues. MIF signalling was strongly associated with DCs in cycling, but more strongly associated with monocytes in repair, and with macrophages in remodelling tissues. Notably, FN1 and SPP1 signalling was strongly associated with monocytes in all conditions.

Due to their strength of association with top signalling pathways, monocytes, macrophages and neutrophils were extracted for further communication analyses (Figure 5F). Pairwise comparisons between experimental conditions (cycling vs repair, cycling vs remodelling, repair vs remodelling) revealed changes in THBS1, SPP1, MIF, CCL and CXCL signalling pathways in monocytes, macrophages and neutrophils relative to phases of endometrial repair and remodelling (Figure 5F). The greatest interaction strength for incoming and outgoing signals in monocytes was associated with THBS1 in repair and SPP1 in remodelling. Macrophage signalling also differed by condition with greatest signalling strength for THBS1 associated with repair while MIF and SPP1 were associated with remodelling. In each instance these signalling pathways were associated with a large differential signal strength for both incoming and outgoing interactions. Neutrophil signalling was also associated with THBS1 and SPP1 in repair and remodelling respectively. CCL signalling strength was reduced in monocytes and macrophages but increased in neutrophils in remodelling. CXCL signalling was strongly associated with neutrophils and both outgoing and incoming signalling strength was upregulated during repair (24 hr) and remodelling (48 hr). Monocytes, macrophages and neutrophils were strongly associated with THBS1 signalling during endometrial repair and SPP1 and MIF signalling during remodelling. Chemokine signalling was distinctly associated with neutrophils during remodelling consistent with roles in immune cell crosstalk and recruitment at this time point.

## Discussion

In this study, we combined a mouse model of menstruation with comprehensive immune cell profiling to characterise the contribution of distinct immune subsets to endometrial repair. We found that uterine immune cell composition was dynamic, with pronounced temporal and spatial changes in monocytes, macrophages and neutrophils which were associated with regions of tissue repair and remodelling. These data suggest that monocyte and neutrophil infiltration is a key feature of endometrial repair, typical of early inflammation responses observed in other tissue contexts[33, 34]. Neutrophils, typically associated with acute inflammatory responses, were abundant during tissue repair as well as remodelling, while macrophages, which regulate tissue repair processes, were relatively decreased during active repair. Myeloid cells were enriched for both pro- and anti-inflammatory signalling pathways, highlighting the discrete balance of immune cell functions that support this unique repair process. Utilising scRNAseq data we were able to identify key signalling pathways associated with myeloid cells during menstruation which included THBS1 and SPP1 as important mediators of endometrial repair. Collectively, these data provide new insights into the immunology of menstruation and highlight specific spatiotemporal roles for myeloid cells during endometrial repair.

We found that monocytes and macrophages were transcriptionally similar with overlapping expression of markers such as *Csf1r*, *Cd68*, *Msr1* and *Fn1*. However, *Inha* and *Cxl3* expression was distinct to monocytes consistent with potential roles in regulating migration and adhesion of inflammatory cells to sites of tissue repair [35]. GOBP analysis highlighted enrichment for genes associated with wound healing and endoplasmic reticulum stress. Monocytes expressed apoptosis- and phagocytosis-associated genes as well as a subset of wound healing genes, such as *Pdpn* and *Cav1*, that were distinct from those expressed by macrophages and neutrophils. Specifically, *Cav1* expression has been shown to increase during monocyte adhesion and drives monocyte-to-macrophage differentiation [36].

CD14+ monocytes were located in both the decidualised tissue and underlying endometrium and were detected in close association with macrophages and neutrophils. Cell chat analysis identified monocytes as major drivers of intercellular communication in the uterus with the highest incoming and outgoing signalling strength amongst immune cells, followed by macrophages and neutrophils. Monocytes were associated with signalling via CCL, MIF, SPP1 and THBS and were the predominant outgoing signal source for these signalling pathways. Signalling via THBS and SPP1 was strongly associated with monocytes from repair and remodelling tissues, respectively.

SPP1 (Secreted Phosphoprotein 1, also known as Osteopontin) plays multiple roles in tissue repair and is reported to modulate inflammation and tissue remodelling. SPP1 mediates cell-to-cell and cell-to-matrix interactions, via integrins and CD44, which can promote macrophage migration and effector functions[37]. SPP1 also increases expression of S100A9, IL-1β, and chemokines like CCL2, CCL3 and CCL4 which skews monocytes toward a pro-inflammatory phenotype[38]. Notably, these genes were enriched in the transcriptome of uterine monocytes during menstruation in the current study. Thrombospondin 1 (TSP-1) promotes cell migration and tissue remodelling via TGF-β and integrin signalling which may support tissue repair in the endometrium [39] [40]. However, it can also inhibit repair by suppressing angiogenesis, a crucial component of endometrial repair [41]. TSP-1 signalling was enriched in monocytes from repair tissues, but the signal strength was relatively decreased in remodelling tissues, when vascular remodelling is more prominent, highlighting the discrete and dynamic contributions of different inflammatory cells and mediators to endometrial repair during menstruation.

We have previously shown that distinct subsets of macrophages associate with regions endometrial repair during menstruation [16] but the function/phenotype of uterine macrophages has not been elucidated. In the current study, we found that macrophages were enriched for signalling pathways important for tissue repair and remodelling. This included complement pathway which has roles in clearing cellular debris, recruiting immune cells to sites of injury, releasing growth factors and regulating blood clotting in tissue repair processes. Differential gene expression analysis highlighted that uterine macrophages distinctly expressed key components of the complement pathway, including *C1qa*, which is a marker of tissue resident macrophages in other contexts [42]. Macrophages were also enriched for genes involved in galectin signalling indicative of a pro-resolving phenotype [43]. Similar to monocytes, macrophages were also associated with MIF and FN1 signalling networks which were key pathways of cell-cell communication. The transcriptomic similarities between macrophages and monocytes are likely representative of linked differentiation trajectories within uterine tissues. Although the ontogeny and kinetics of monocytes and macrophages in the uterus are yet to be fully defined, studies using tissue explants, or organ transplants, suggest the uterine macrophage niche is maintained via monocyte-driven replenishment [44, 45]. This is consistent with the compositional changes in macrophages and monocytes described in the current study, where monocytes are predominant during repair, but macrophages, likely monocyte-derived, are more abundant than monocytes during remodelling.

ScRNAseq data analysis confirmed that neutrophils were a major source of chemokines as they were enriched for both CCL and CXCL-mediated signalling pathways, indicating regulation of immune cell recruitment within the repairing endometrium. Neutrophils were associated with pro-inflammatory signalling via TNF pathways, which, interestingly, was prominent during remodelling rather than repair. While TNF is a classic pro-inflammatory mediator, Annexin1 signalling was also enriched in neutrophils at this timepoint. Annexin1 has anti-inflammatory functions by downregulating neutrophil activation [46, 47]. The neutrophil transcriptome profile was consistently associated with both pro- and anti-inflammatory pathways which may suggest neutrophil phenotype is temporally altered in the uterine tissue environment across repair and remodelling, with a switch to anti-inflammatory functions during the latter phase. Assessing differentially expressed genes in neutrophils highlighted that in addition to classic neutrophils markers such as *S100a8*, *S100a9* and *Cxcl2*, uterine neutrophils also expressed *Dusp1* and *Acod1* indicating an anti-inflammatory phenotype. Notably, Dusp1 (Dual Specificity Phosphatase 1) has key roles in resolution of inflammation by deactivating pro-inflammatory mitogen-activated protein kinase (MAPK)-dependent pathways and influencing production of neutrophil chemoattractants like Cxcl1[48]. This may reflect a functional requirement for restraining classic effector functions of neutrophils within the uterus in the context of endometrial repair. Expression of Acod1 (Aconitate Decarboxylase 1) in neutrophils regulates inflammation via production of itaconate. This shift in immunometabolism is associated with anti-inflammatory protein production, protecting cells from oxidative stress as well as regulation of NET formation and neutrophil migration [49, 50]. Thus, although neutrophils are abundant during menstruation, their transcriptomic phenotype supports roles in coordinating immune responses via chemokine secretion and promoting resolution of inflammation through suppression of pro-inflammatory signalling pathways. Transcriptome profiling in the current study supports reparative functions for neutrophils during menstruation consistent with emerging roles for neutrophils in contributing to tissue repair in other contexts [51–53]. Indeed, Kaitu’u-Lino et al. previously demonstrated the importance of monocytes and macrophages to endometrial breakdown and repair using antibody-based depletion (RB6-8C5) of Gr-1 positive cells in a mouse model of menstruation [54].

Our findings using a mouse model of simulated menstruation extend and complement those from analysis of human endometrial tissues. Armstrong et al., investigated mRNA expression levels of inflammatory cytokines and chemokines during menstruation in the human endometrium and found increased expression of CCL2, CXCL8, IL6 and TNF during the menstrual phase of the cycle[8]. Importantly, the current study provides much needed resolution to the specific cellular sources of these mediators which may be beneficial to translational studies aiming to understand the pathophysiology of menstrual dysfunction in reproductive health disorders. Dysregulated inflammatory responses are a common feature of reproductive health disorders such as heavy menstrual bleeding (HMB) and endometriosis, with changes to macrophage and neutrophil function reported[55]. While studies characterising specific phenotypes of immune cell subsets during menstruation are lacking, changes in expression to key mediators described in the current study have been reported. For example, TSP-1 expression is reported to be decreased in endometrial tissues from women with HMB [41] while overexpression of SPP1 has been reported in macrophages from women with endometriosis[56]. Further research will be needed to better characterise immune dysfunction in human reproductive disorders, but the comprehensive phenotyping performed in the current study provides tractable targets for further translational investigation.

## Conclusion

Here, we have comprehensively profiled immune cells using a combination of scRNAseq, flow cytometry and multiplex immunohistochemistry and identified that monocytes, macrophage and neutrophils are spatially and temporally regulated to support endometrial repair. These data provide a more complete understanding of how different immune cell populations contribute to endometrial tissue repair and identify key cell-specific mediators and pathways. Future studies should assess the importance of myeloid cells to endometrial repair through lineage tracing, cell ablation and comparison to human datasets. Improved understanding of endometrial repair will, in turn, support identification of potential pathways that are dysregulated in reproductive health disorders in women.

## General

We are sincerely grateful to the members of the SURF imaging suite at the University of Edinburgh and the Veterinary and other staff in Biological Services for their assistance. Flow cytometry data were generated with support from the IRR Flow Cytometry and cell sorting facility, University of Edinburgh.

For the purpose of open access, the author has applied a Creative Commons Attribution (CC BY) licence to any Author Accepted Manuscript version arising from this submission.

## Author contributions

Conceptualization: DAG

Methodology: RJA, PMK, HL, SG, IS, DAG

Analysis: RJA, PMK, HL, SG, IS, DAG

Funding acquisition: DAG

Writing – original draft: RJA, PMK, IS, DAG

Writing – review & editing: RJA, PMK, PTKS, IS, DAG

## Data and materials

All data are available in the main text or the supplementary materials

## Supplementary data

**Supplementary figure 1:**
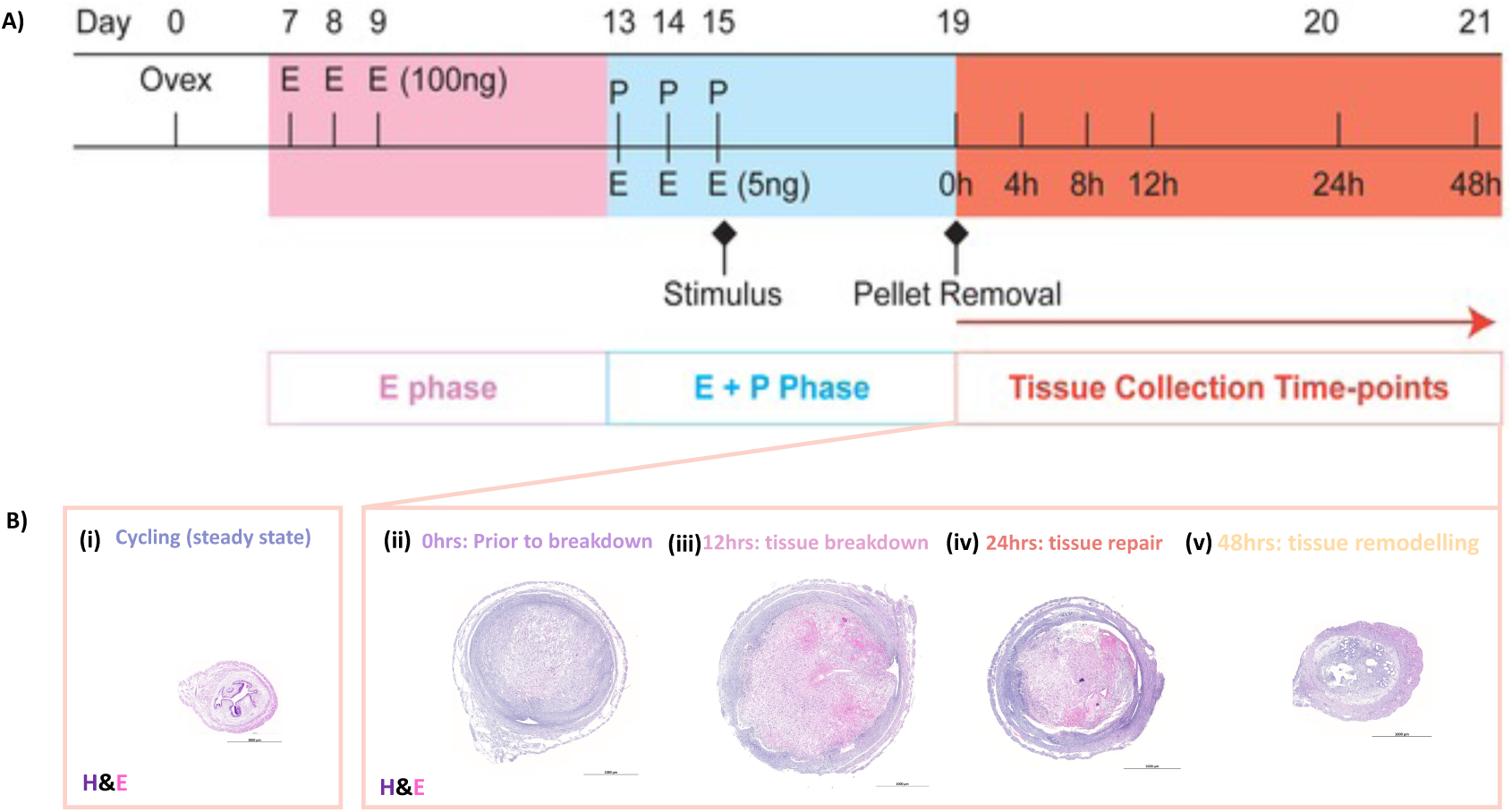
Mouse model of simulated menstruation characterised by endometrial tissue breakdown and repair. **(A)** Schematic detailing the timeline of surgical procedures and hormonal stimulation used to induce a menstruation-like event in mice. **(B)** Haematoxylin and eosin (H&E) staining of uterine tissue sections demonstrate macro- and microscopic tissue architecture in **(i)** steady state mouse uterus and mouse uterus at various time points of tissue breakdown and repair: **(ii)** 0hrs, prior to breakdown; **(iii)** 12hrs, tissue breakdown; **(iv)** 24hrs, endometrial repair; **(v)** 48hrs, endometrial remodelling (time post progesterone withdrawal; scale bar=1000µm).

**Supplementary Table 1:**
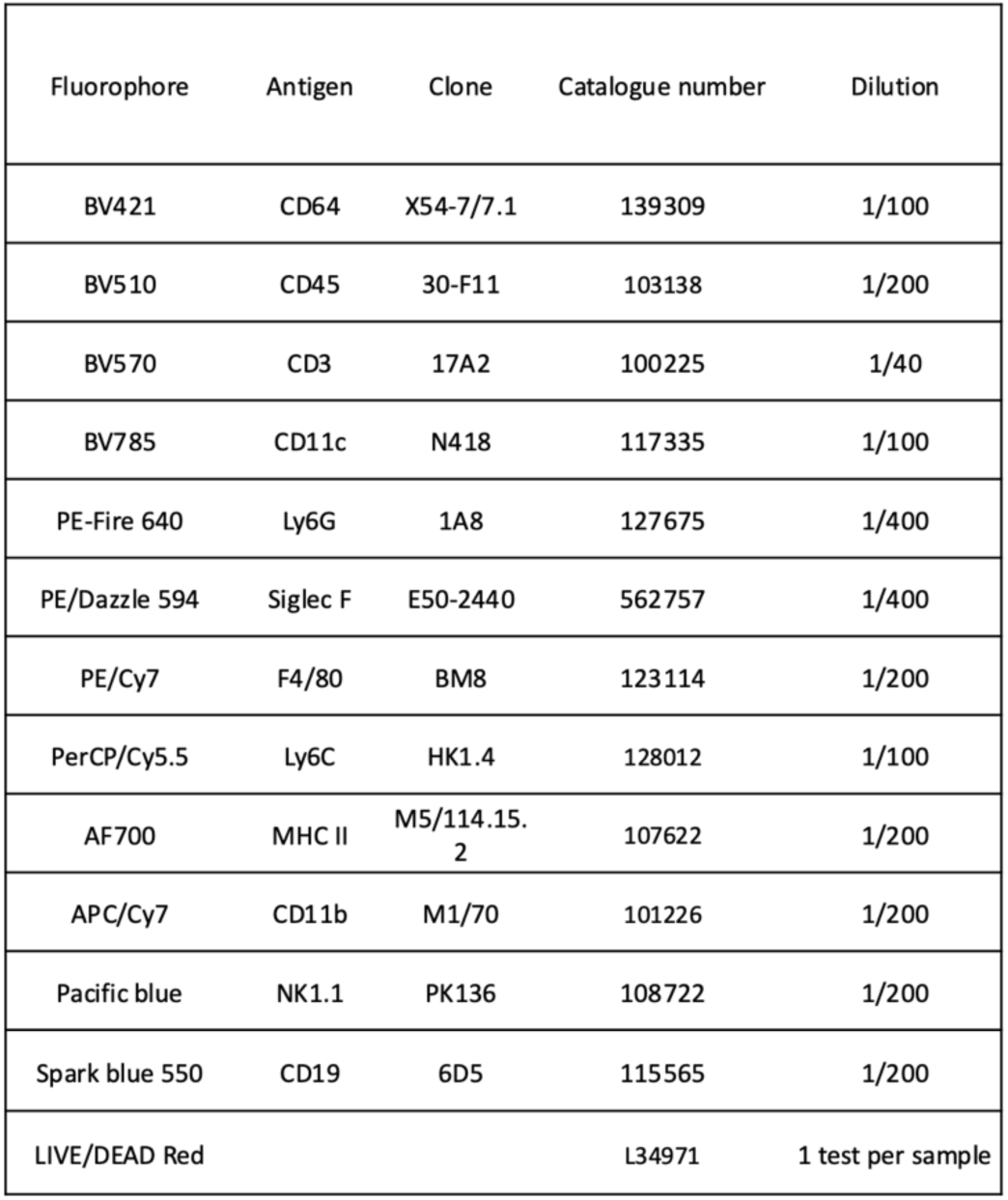
Flow cytometry antibody panel utilised to characterise uterine immune cell dynamics.

| Fluorophore | Antigen | Clone | Catalogue number | Dilution |
| --- | --- | --- | --- | --- |
| BV421 | CD64 | X54-7/7.1 | 139309 | 1/100 |
| BV510 | CD45 | 30-F11 | 103138 | 1/200 |
| BV570 | CD3 | 17A2 | 100225 | 1/40 |
| BV785 | CD11c | N418 | 117335 | 1/100 |
| PE-Fire 640 | Ly6G | 1A8 | 127675 | 1/400 |
| PE/Dazzle 594 | Siglec F | E50-2440 | 562757 | 1/400 |
| PE/Cy7 | F4/80 | BM8 | 123114 | 1/200 |
| PerCP/Cy5.5 | Ly6C | HK1.4 | 128012 | 1/100 |
| AF700 | MHC II | M5/114.15.<br>2 | 107622 | 1/200 |
| APC/Cy7 | CD11b | M1/70 | 101226 | 1/200 |
| Pacific blue | NK1.1 | PK136 | 108722 | 1/200 |
| Spark blue 550 | CD19 | 6D5 | 115565 | 1/200 |
| LIVE/DEAD Red |  |  | L34971 | 1 test per sample |

**Supplementary figure 2:**
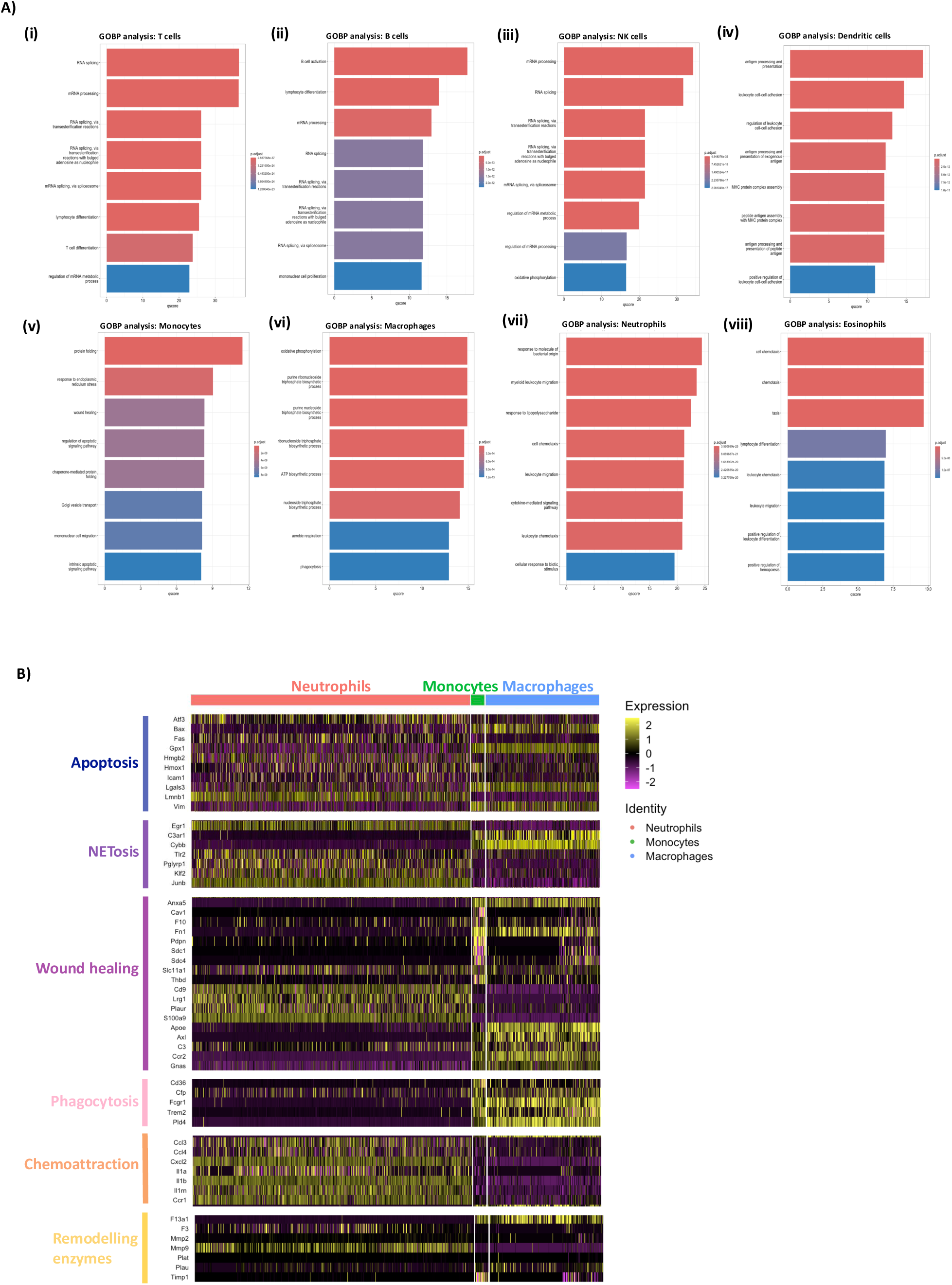
Gene Ontology (GO) analysis reveals biological processes (BP) associated with the transcriptome of each immune cell cluster. **(A)** Bar plot: Unbiased analysis of GOBP terms associated with the genetic signature of each immune cell cluster: **(i)** T cells **(ii)** B cells **(iii)** NK cells **(iv)** Dendritic cells **(v)** Monocytes **(vi)** Macrophages **(vii)** Neutrophils **(viii)** Eosinophils. Bar size= qscore: -log(p.adjust) ; bar colour= P.adjust value. **(B)** Heatmap (yellow, high; purple, low): Biased analysis of gene signatures (GSEA gene sets) associated with known wound tissue repair GO terms by neutrophil, monocyte and macrophage clusters: apoptosis, NETosis, wound healing, phagocytosis, chemoattraction, remodelling enzymes. Top axis is colour coded and named by immune cell type.

**Supplementary figure 3:**
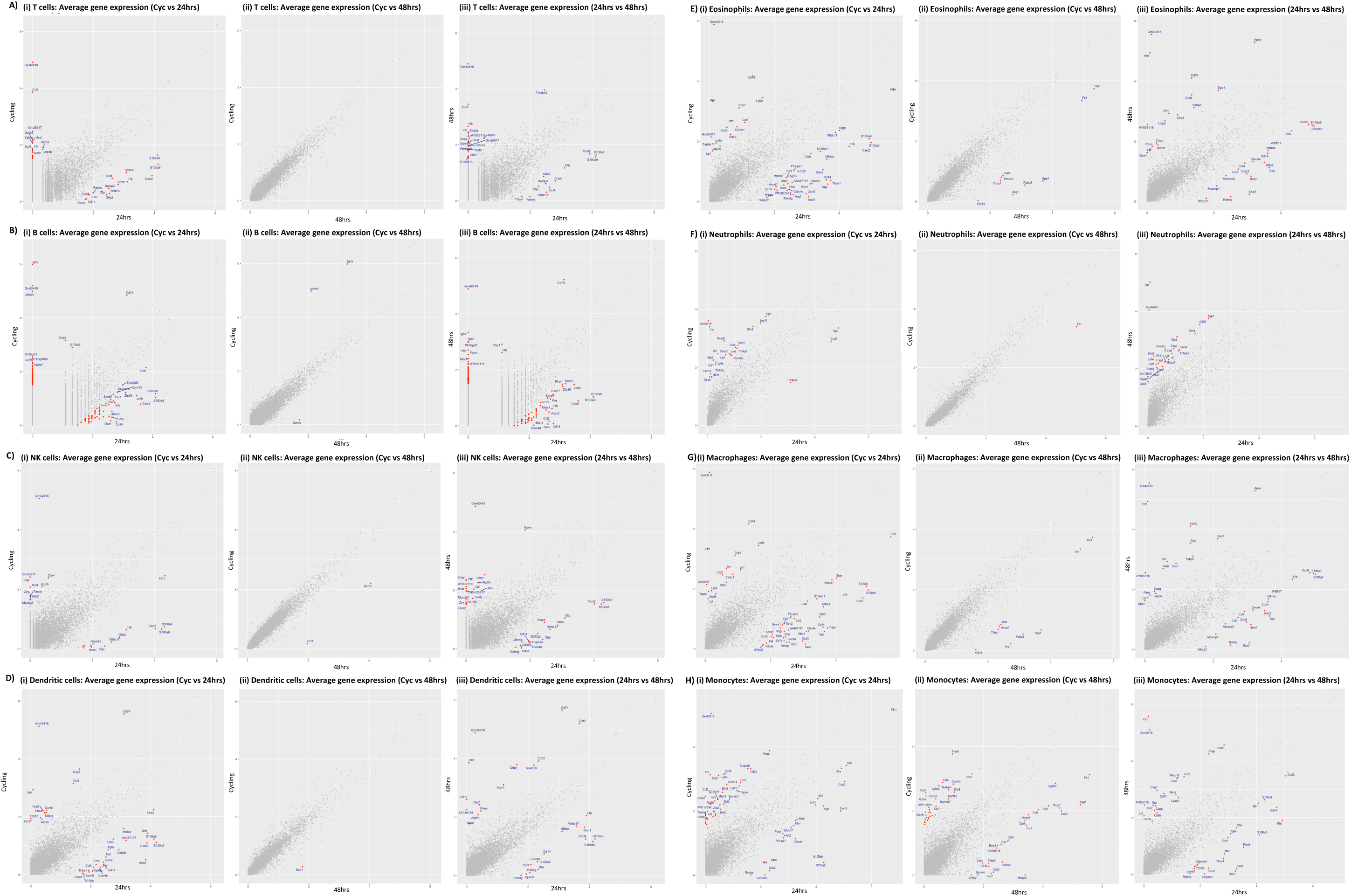
Average gene expression analysis for each immune cell cluster between the 3 experimental conditions. Scatter plot: Average gene expression analysis for each immune cell cluster **(A)** T cells **(B)** B cells **(C)** NK cells **(D)** Dendritic cells **(E)** Monocytes **(F)** Macrophages **(G)** Neutrophils **(H)** Eosinophils between different experimental conditions: **(i)** Cycling vs 24 hr **(ii)** Cycling vs 48 hr **(iii)** 24 hr vs 48 hr.

**Supplementary figure 4:**
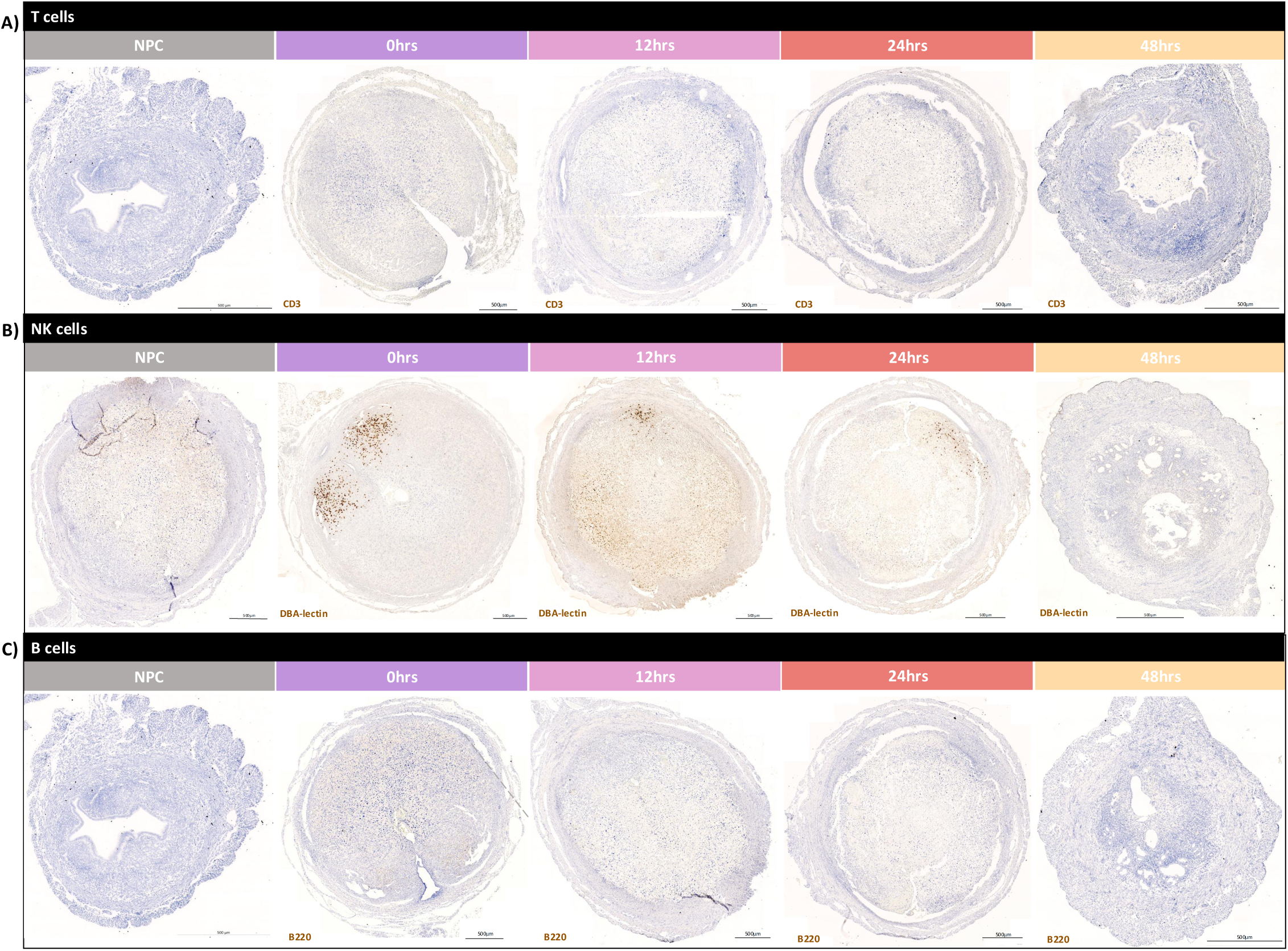
Immunohistochemistry analysis of uterine lymphocyte dynamics during endometrial repair using a mouse model of simulated menstruation. Immunohistochemistry: detection of **(A)** CD3+ T cells, **(B)** DBA-lectin+ NK cells and **(C)** B220+ B cells in uterine tissue sections at 0hr (prior to breakdown, n=4), 12hr (tissue breakdown, n=4-6), 24hr (tissue repair, n=4-5) and 48hr (tissue remodelling, n=4-6) hr after progesterone withdrawal (representative images, scale bar=500µm).

**Supplementary figure 5:**
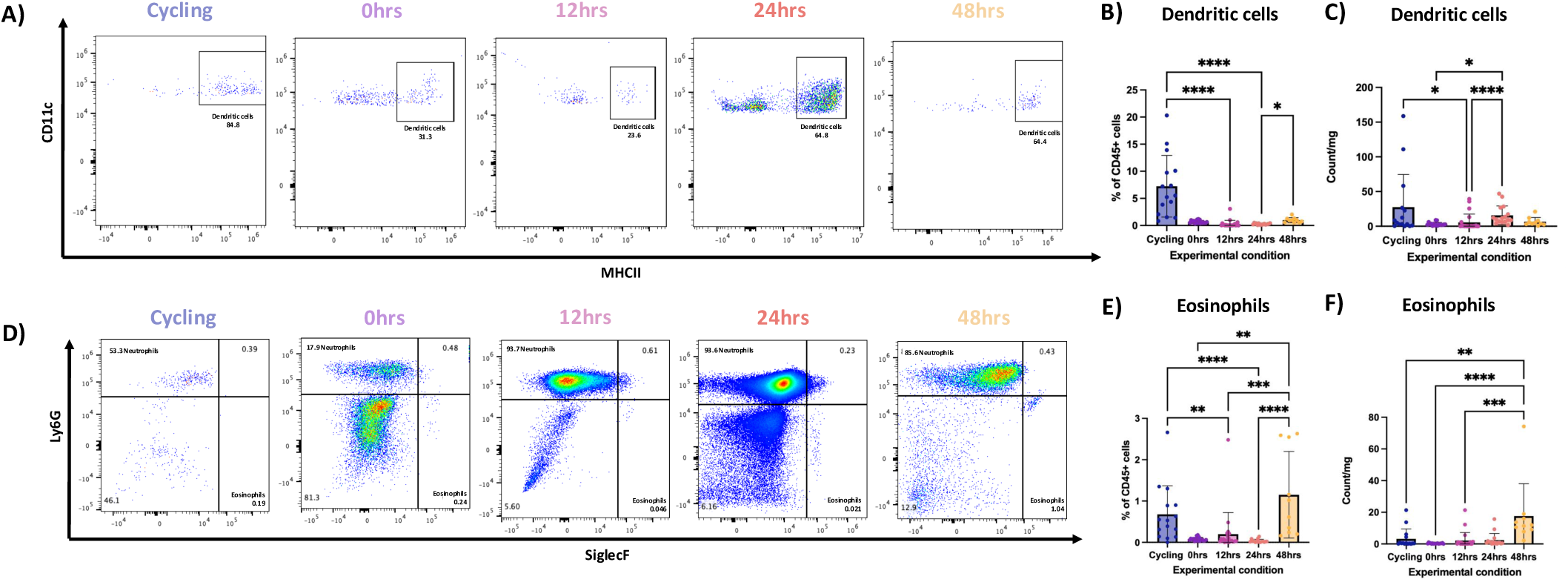
Flow cytometry analysis of dendritic cells and eosinophils in uterine tissues endometrial repair using a mouse model of simulated menstruation. **(A)** Population restricted FC analysis to identify CD45+ CD3- CD19- NK1.1- Ly6G- CD64- CD11c+ MHCII+ dendritic cells (DCs) in uterine tissue digests. **(B)** Bar plot, FC quantification: Relative abundance of DCs calculated as a percentage of total CD45+ cells per uterine horn. **(C)** Bar plot, FC quantification: Absolute counts of DCs per uterine horn. **(D)** Population restricted FC analysis to CD45+ CD3- CD19- NK1.1- CD11b+ Ly6G- SiglecF+ eosinophils in uterine tissue digests. **(E)** Bar plot, FC quantification: Relative abundance of eosinophils calculated as a percentage of total CD45+ cells per uterine horn. **(F)** Bar plot, FC quantification: Absolute counts of eosinophils per uterine horn. (FC analysis groups: control (n=15), 0hr (prior to breakdown, n=17), 12hr (tissue breakdown, n=22), 24hr (tissue repair, n=16) and 48hr (tissue remodelling, n=10) after progesterone withdrawal; statistical comparisons were made using Kruskal-Wallis tests with multiple comparisons. and * p< 0.05, ** p < 0.01, *** p < 0.001, **** p < 0.0001).

**Supplementary figure 6:**
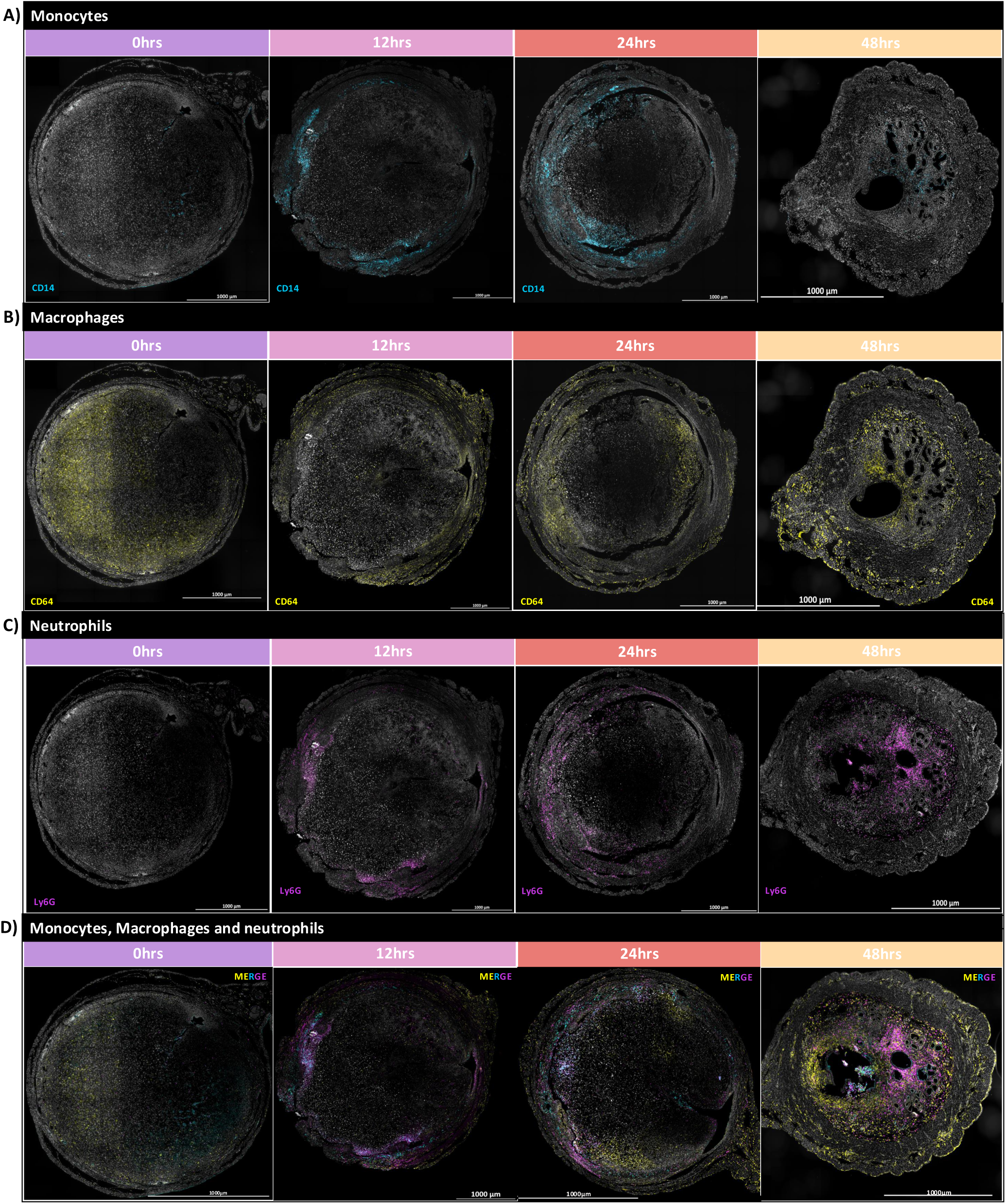
Immunohistochemistry analysis of myeloid cell dynamics during endometrial repair using a mouse model of simulated menstruation. Immunohistochemical detection of **(A)** CD14+ monocytes (cyan), **(B)** CD64+ macrophages (yellow) and **(C)** Ly6G+ neutrophils (magenta) in uterine tissue sections at 0hr (prior to breakdown, n=4), 12hr (tissue breakdown, n=5), 24hr (tissue repair, n=5) and 48hr (tissue remodelling, n=3) following progesterone withdrawal (representative images, scale bar=1000µm). (D) Merged channel representative images (scale bar=1000µm).

## Appendix

### Abbreviations

#### Abbreviation Full term

Acod1: Aconitate decarboxylase 1
Adgre1: Adhesion G protein–coupled receptor E1 (F4/80)
Anxa5: Annexin A5
Apoe: Apolipoprotein E
APP: Amyloid precursor protein
AWERB: Animal Welfare and Ethical Review Body
Axl: AXL receptor tyrosine kinase
Cav1: Caveolin 1
C3: Complement component 3
Ccl2: C–C motif chemokine ligand 2
Ccl3: C–C motif chemokine ligand 3
Ccl4: C–C motif chemokine ligand 4
Ccl5: C–C motif chemokine ligand 5
Ccl6: C–C motif chemokine ligand 6
Ccl9: C–C motif chemokine ligand 9
CCL: C–C motif chemokine ligand
CCR2: C–C motif chemokine receptor 2
Cd3e: CD3 epsilon chain
CD3: Cluster of differentiation 3
CD9: Cluster of differentiation 9
CD11b: Integrin alpha M
CD11c: Integrin alpha X
CD14: Cluster of differentiation 14
CD19: Cluster of differentiation 19
CD45: Protein tyrosine phosphatase receptor type C
CD64: Fc gamma receptor I
Cd68: Cluster of differentiation 68
Cdc1: Conventional dendritic cell 1
cDNA: Complementary DNA
Clec9a: C-type lectin domain family 9 member A
Clec10: C-type lectin domain family 10
Csf1r: Colony stimulating factor 1 receptor
CXCL: C-X-C motif chemokine ligand
Cxcl1: C-X-C motif chemokine ligand 1
Cxcl3: C-X-C motif chemokine ligand 3
Cxcr2: C-X-C motif chemokine receptor 2
CXCL8: C-X-C motif chemokine ligand 8
DCs: Dendritic cells
DEGs: Differentially expressed genes
Dusp1: Dual specificity phosphatase 1
E2: Estradiol
EDTA: Ethylenediaminetetraacetic acid
F10: Coagulation factor X
F4/80: Macrophage marker (Adgre1)
FACS: Fluorescence-activated cell sorting
FC: Flow cytometry
FFPE: Formalin-fixed paraffin-embedded
FN1: Fibronectin 1
Fn1: Fibronectin 1
FSC-A: Forward scatter area
FSC-H: Forward scatter height
GEMs: Gel bead-in-Emulsions
GO: Gene ontology
GOBP: Gene ontology biological processes
Gata3: GATA binding protein 3
Gnas: GNAS complex locus
Gzmb: Granzyme B
H2-Ab1: MHC class II antigen A, beta 1
Hic1: Hypermethylated in cancer 1
HMB: Human menopausal blood
HRP: Horseradish peroxidase
Ighd: Immunoglobulin heavy constant delta
IL-1β: Interleukin 1 beta
IL-6: Interleukin 6
Inha: Inhibin alpha
ISG15: Interferon-stimulated gene 15
IFIT1: Interferon-induced protein with tetratricopeptide repeats 1
IFIT3: Interferon-induced protein with tetratricopeptide repeats 3
Klrb1c: Killer cell lectin-like receptor B1C
Lrg1: Leucine-rich alpha-2-glycoprotein 1
Ly6C: Lymphocyte antigen 6 complex, locus C
Ly6G: Lymphocyte antigen 6 complex, locus G
MAPK: Mitogen-activated protein kinase
Mertk: MER proto-oncogene tyrosine kinase
MHC II: Major histocompatibility complex class II
MIF: Macrophage migration inhibitory factor
MMP: Matrix metalloproteinase
Msr1: Macrophage scavenger receptor 1
NET: Neutrophil extracellular trap
NK1.1: Natural killer cell marker
Ncr1: Natural cytotoxicity triggering receptor 1
P4: Progesterone
PCA: Principal component analysis
Pdpn: Podoplanin
PBS: Phosphate-buffered saline
Plaur: Plasminogen activator, urokinase receptor
QC: Quality control
RNA: Ribonucleic acid
Rsad2: Radical S-adenosyl methionine domain containing 2
S100a8: S100 calcium-binding protein A8
S100a9: S100 calcium-binding protein A9
ScRNAseq: Single-cell RNA sequencing
Sdc1: Syndecan 1
Sdc4: Syndecan 4
Sell: L-selectin
SiglecF: Sialic acid-binding Ig-like lectin F
Slc11a1: Solute carrier family 11 member 1
SPP1: Secreted phosphoprotein 1
SSC-A: Side scatter area
SSC-H: Side scatter height
TGF-β: Transforming growth factor beta
TGFBI: Transforming growth factor beta-induced
THBS1: Thrombospondin 1
Thbd: Thrombomodulin
Tmem123: Transmembrane protein 123
TNF-α: Tumour necrosis factor alpha
TPM: Transcripts per million
TSP-1: Thrombospondin-1
UMAP: Uniform Manifold Approximation and Projection
uNKs: Uterine natural killer cells
VEGF: Vascular endothelial growth factor
Xcr1: X-C motif chemokine receptor 1

